# Predictive learning shapes the representational geometry of the human brain

**DOI:** 10.1101/2024.03.07.583842

**Authors:** Antonino Greco, Julia Moser, Hubert Preissl, Markus Siegel

## Abstract

Predictive coding theories propose that the brain constantly updates its internal models of the world to minimize prediction errors and optimize sensory processing. However, the neural mechanisms that link the encoding of prediction errors and optimization of sensory representations remain unclear. Here, we provide direct evidence how predictive learning shapes the representational geometry of the human brain. We recorded magnetoencephalography (MEG) in human participants listening to acoustic sequences with different levels of regularity. Representational similarity analysis revealed how, through learning, the brain aligned its representational geometry to match the statistical structure of the sensory inputs, by clustering the representations of temporally contiguous and predictable stimuli. Crucially, we found that in sensory areas the magnitude of the representational shift correlated with the encoding strength of prediction errors. Furthermore, using partial information decomposition we found that, prediction errors were processed by a synergistic network of high-level associative and sensory areas. Importantly, the strength of synergistic encoding of precition errors predicted the magnitude of representational alignment during learning. Our findings provide evidence that large-scale neural interactions engaged in predictive processing modulate the representational content of sensory areas, which may enhance the efficiency of perceptual processing in response to the statistical regularities of the environment.

## Introduction

Living organisms must adapt to ever-changing complex environments. To accomplish this, it is advantageous to anticipate environmental changes that posit threats or opportunities to survival. For this reason, the brain is able to detect and extract statistical regularities in sensory inputs, an ability that has been referred to as statistical learning and which humans are especially capable of (1–3).

Predictive coding theory provides a framework to explain how regularities are extracted from sensory inputs and how they are used to optimally predict future outcomes (4–6). This framework generally assumes that the brain possesses a generative internal model of the environment (latent variables that cause the sensory observations), and updates the model by computing the prediction error between its probabilistic predictions and the sensory data (7–9). These predictive mechanisms are thought to give rise to increased neural activity following an unexpected sensory input (10, 11) or decreased neural responses when an input is expected (12, 13). In the auditory domain, the oddball paradigm has been widely adopted to probe the ability of the brain to track statistical regularities (14). A rare deviant tone presented within a sequence of regular tones elicits an event-related response, the so called mismatch negativity (MMN) (15). Under the predictive coding framework, this response is interpreted as a neural signature of the prediction error between the expectation of the generative model (regular tone) and the sensory data (deviant tone) (16, 17). Furthermore, recent studies on auditory statistical learning that used more complex patterns of sound sequences showed that cortical responses do not only reflect violations of sensory predictions at a local tone level but also on the global sequence level (18–22).

In sum, a large body of evidence has provided insights into cortical signals compatible with the encoding of prediction errors. In contrast, little is known how such signals are used to update the brains internal generative model. Studies on perceptual learning suggest plasticity of sensory representations even in low-level sensory regions to optimize sensory processing (23–25). Furthermore, studies on statistical learning show that the similarity of neuronal representations of sensory stimuli reflects the learned statistical dependencies between these stimuli (26–28). This suggests that statistical learning may shape the geometry of neural representations to match the geometry of sensory inputs. However, the neural mechanisms underlying this learning remain unclear. If prediction error signals are used to update neuronal representations these two phenomena should be linked, i.e. the neuronal encoding of predictions errors should be correlated with the updating of neuronal repre sentations. However, so far evidence to support this fundamental link between prediction errors and the updating of neural representations is missing. Here, we sought to establish this link.

We performed magnetoencephalography (MEG) recordings in human participants passively listening to acoustic tone triplet sequences with low or high regularity. Representational similarity analysis (29) revealed that, through learning, the brain aligned its representational geometry to match the statistical structure of the sequences. We employed computational modelling to derive neural signals encoding prediction error trajectories (30–32). We found that the strength of prediction error encoding indeed predicted the magnitude of the alignment of sensory representations through learning. Furthermore, based on Partial Information Decomposition (33) we found that brain regions that showed representational alignment also engaged in a synergistic encoding of prediction errors. Our findings suggest that in the human brain large-scale neural interactions engaged in predictive processing modulate the sensory representational geometry in response to the statistical regularities of the environment.

## Results

We recorded MEG in 24 human participants that passively listed to two sequences of 12 acoustic tones (34) (Fig. 1A). Each tone had a duration of 300 ms followed by a 33 ms silent gap. For the sequence construction, tones were grouped into triplets (1 s triplet duration) where the tones in a triplet never spanned more than one octave (Fig. 1B). Both sequences consisted of a total of 800 triplets. One sequence, which we are referring to as the high regularity (HR) condition, consisted of only four different types of triplets, while in the other sequence, which we are referring to as the low regularity (LR) condition, the order of tones inside a triplet was changing throughout the sequence (Fig. 1C). The different regularities of the two sequences are reflected in their distinct transition matrices between consecutive tones (Fig. 1C right). We source-reconstructed neural activity throughout the brain from the MEG data using the Desikan-Killany parcellation scheme (35) and beamforming (36). Both tone sequences evoked robust neural responses that peaked about 60 ms after each tone onset in bilateral auditory cortices (Fig. 1D). We applied multivariate decoding and representational similarity analysis (RSA) (29) on the source-level brain activity to investigate the neural representation of different tones and to test if this representation changed during learning (Fig. 2). We quantified the distance between neural population responses to different tones using the cross-validated Mahalanobis distance (cvMD) (37, 38) (Fig. 2A). This yielded representational dissimilarity matrices (RDMs) that quantified the distance of neural representations for all pairs of tones. Importantly, we performed this analysis temporally resolved relative to each tone presentation and separately in five consecutive blocks of trials throughout each sequence. This allowed us to resolve the temporal dynamics of neuronal tone representation on a fast timescale in response to each tone and on a slow timescale throughout learning. As a first step, we averaged all RDMs across tone pairs and blocks. We found that tones were well decodable for both sequences, with peak decoding performance around 100 ms post tone onset (Fig. 2B, p < 0.0001 cluster-corrected; peak Cohen’s d = 1.46). We next ordered RDM entries such that values near and off the diagonal represented representational distances of tones within and between triplets in the HR sequence, respectively. Then, we separately quantified neural distances of tones within (Fig. 2C left) and between triplets (Fig. 2C middle) across time and blocks. To quantify the representational dynamics during learning, we computed the linear slope of the average representational distance within and between triplets across sequence blocks (Fig. 2C bottom). We predicted that if learning matched the representational geometry of neural representations to the statistical regularity of sequences, then distances of tones within triplets should decrease across blocks more than of tones between triplets and this effect should be specific for the high regularity condition. Indeed, this is what we found.

**Figure 1:**
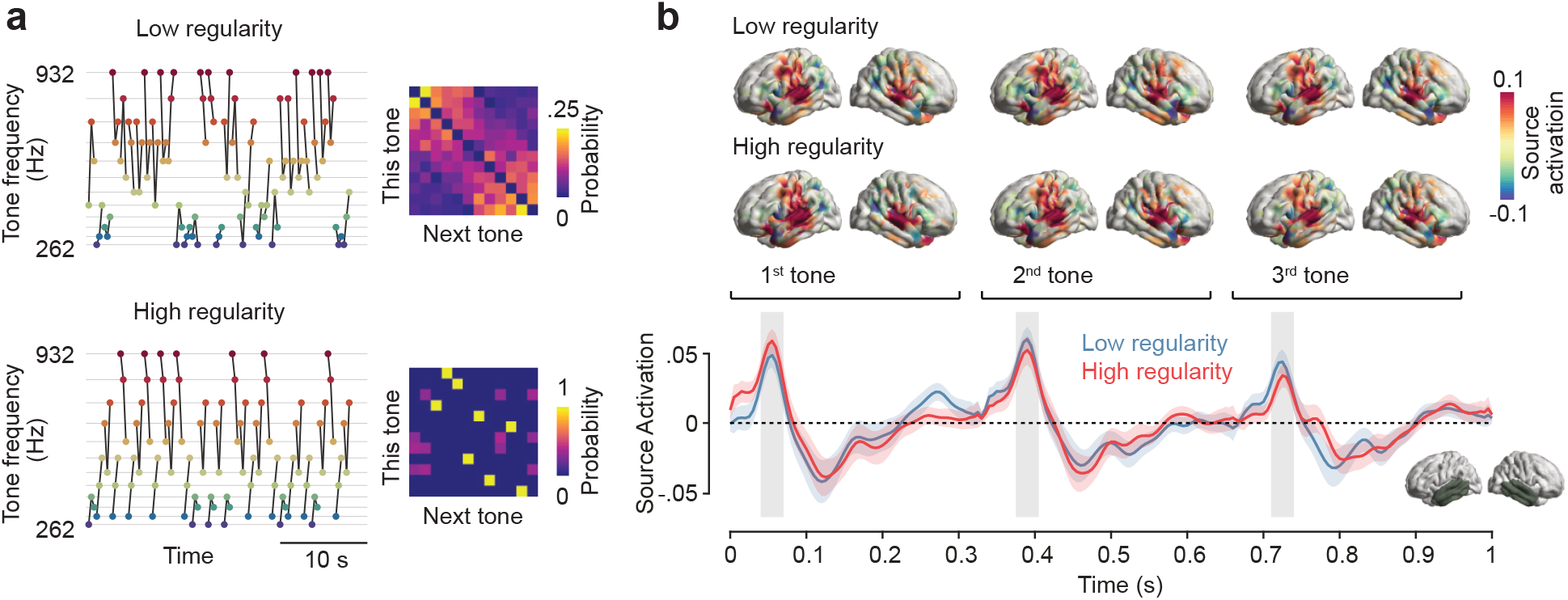
Experimental design and auditory cortical responses. Subjects passively listened to two sequences of 12 acoustic tones. **a**, left: exemplary sections of low and high regularity sequences arranged by their frequency. Grey lines indicate triplets. Right: transition matrices of subsequent tones for both sequences. **b**, top: cortical distribution of evoked activity 50-70 ms post onset of each of the three tones in a triplet. Bottom: source-reconstructed evoked activity in bilateral temporal cortices (bottom right inset) across triplets in the low and high regularity condition. Shaded areas indicate standard error of the mean (SEM).

**Figure 2:**
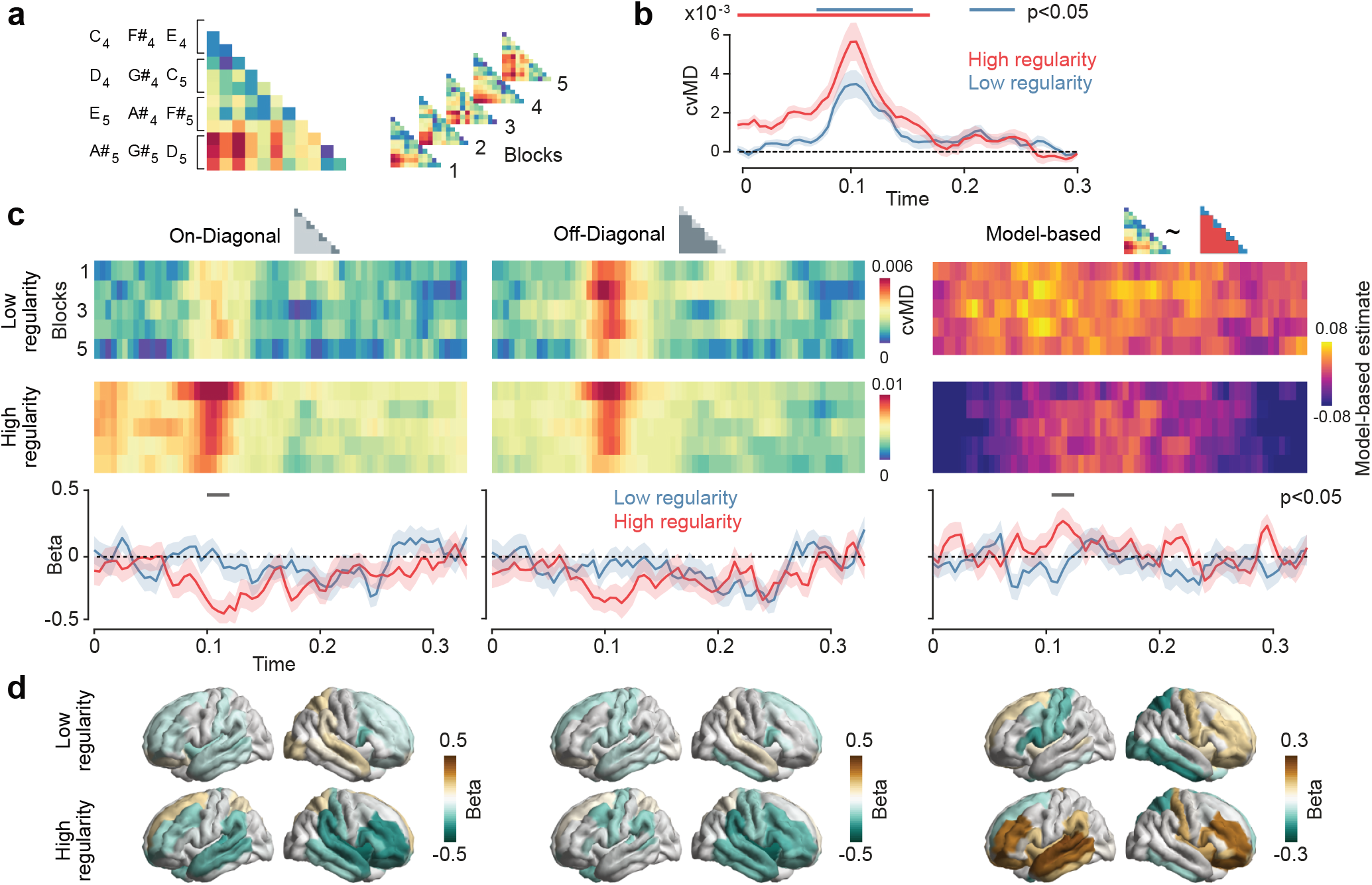
Representational Similarity Analysis (RSA) showing the “representational shift” for the high regularity sequence. **a**, Analysis pipeline. Representational dissimilarity matrices (RDMs) were computed based on the cross-validated Mahalanobis Distance (cvMD) between all pairs of tones and ordered according to the triplet structure in the high regularity sequence. RDMs were computed for 5 consecutive blocks of trials. **b**, Time course of a the average RDM for tones in both sequences. Shaded areas indicate SEM and horizontal lines indicate statistical significance. **c**, Top: on- and off-diagonal cvMD and model-based RSA estimates plotted as a function of time and blocks. Bottom: Regression slopes of cvMD values and RSA coefficients across blocks. Shaded areas indicate SEM and horizontal lines indicate statistical significance. **d**. Searchlight RSA for on- and off-diagonal cvMD values and model-based estimates across the brain in the 110-120 ms time window.

For the high regularity condition, around 120 ms post tone onset, representational distances within triplets decreased across blocks (negative slope) and this decrease was significantly stronger for the high as compared to low regularity condition (Fig. 2C left; p = 0.008 cluster-corrected; d = 0.79). There was also a trend for representational distances to decrease between triplets, but there was no significant difference of slopes between conditions (Fig. 2C right; p > 0.05 cluster-corrected). To directly test our prediction, we then performed a model-based RSA and fitted a theoretical RDM in which the within-triplet distances were lower than between-triplet distances (Fig. 2C). As predicted, around 120 ms the model-fit increased across blocks in the high regularity condition and this increase was significantly stronger than in the low regularity condition (p = 0.038 cluster-corrected, d = 0.84). Which brain regions showed this updating of sensory representations? To address this question, we repeated the analysis in a searchlight fashion across the cortical surface for the time interval from 110 ms to 120 ms post tone onset (Fig. 2D). We found that the decrease of representational distances across blocks within and between triples was strongest in dorsolateral prefrontal and temporal regions (Fig. 2D). Also, the model-based RSA showed that the increase of the model fit across blocks, i.e. the relative decrease of representational distances within triplets, in the high regularity condition peaked in bilateral dorsolateral prefrontal cortices and in left temporal cortex (Fig. 2D). In sum, these findings showed that the brain changed its sensory representations to adaptively match the statistical structure of the sensory inputs. Specifically, in the context with predictable tone triplets, the brain updated the tones’ representations a way that made them more similar between tones belonging to the same triplet as compared to tones belonging to different triplets.

After establishing how statistical learning updated sensory representations, we next focussed on the neural encoding of errors between sensory inputs and input predictions (6, 30). We adopted a computational modelling approach to examine how the brain encoded the prediction error. We used an ideal observer model (Fig. 3A), a perceptron neural network resembling the Rescorla-Wagner model for categorical data (30), that predicted the next tone given the previous one. Inspired by Bayesian models of predictive coding, the model employed a dynamic learning rate that changed as a function of the ideal observer’s uncertainty (39). After fitting the model to each stimulus sequence, the weight matrix captured the actual transition matrix of each sequence (Fig. 3B, compare Fig. 1C). Also, the prediction error and accuracy of the fitted models reflected the statistics of the two sequences with lower prediction error and higher accuracy for the more predictable high regularity condition (Fig. 3B). We then extracted the prediction error trajectories of the model for each condition (Fig. 3B) and tested if prediction errors were encoded by the neural activity using Gaussian Copula Mutual Information (GCMI) (40) (Fig. 3A). We found that the prediction errors were indeed significantly encoded, peaking around 100 ms post tone onset for both high and low regularity sequences (Fig. 3C) (HR: 0-50 ms, p < 0.0001 cluster-corrected, d = 0.78; LR: first cluster 0-40 ms, p = 0.007 cluster-corrected, d = 0.94, second cluster 50-260 ms, p < 0.0001 cluster-corrected, d = 1.72). There was no significant difference of the prediction error encoding between the sequences (p > 0.05 cluster-corrected). Again, we repeated the analysis in a searchlight fashion across the cortical surface to investigate the cortical distribution. Prediction errors were encoded broadly across frontoparietal and temporal areas, with the right temporal cortex showing maximum neural prediction error signals in both sequences (Fig. 3C bottom).

**Figure 3:**
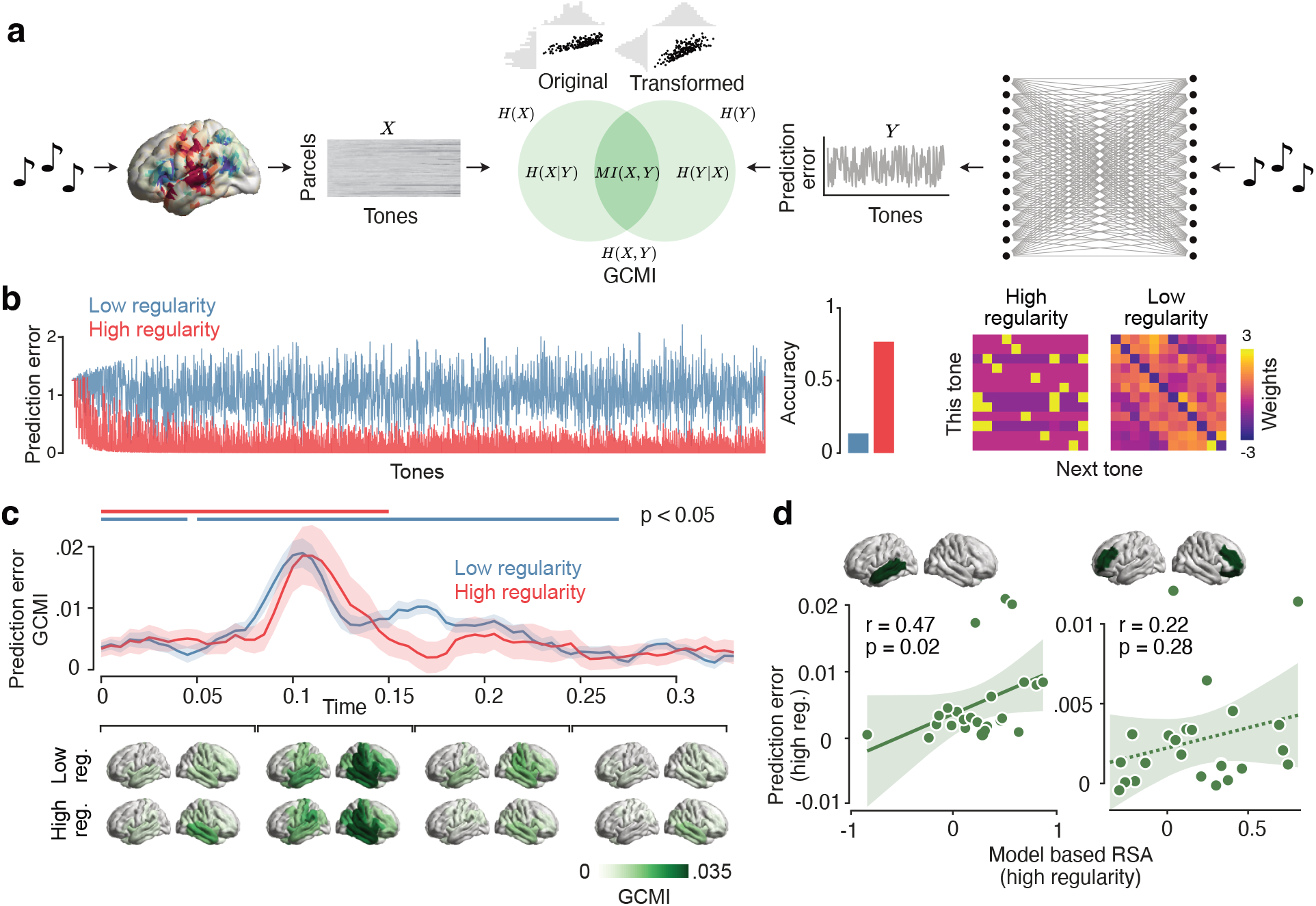
Computational modeling of prediction error trajectories using an ideal observer model. **a**, Analysis framework for fitting prediction error trajectories from the model to each parcel of the brain. Left: For each time point relative to a tone presentation, the brain data is structured as a matrix with parcels and tones. Right: prediction error trajectory extracted from the ideal observer model. Centre: illustration of Gaussian Copula Mutual Information (GCMI). **b**, Left: prediction error trajectories extracted from the model for both sequences. Middle: prediction accuracies of the model. Right: weight matrices of the ideal observer model after training on the full sequences. **c**, top: Time-course of prediction error encoding (GCMI) for both sequences. Shaded areas indicate SEM and horizontal lines indicate statistical significance. Bottom: cortical distribution of prediction error encoding in 4 time windows. **d**, correlation analysis between the representational shift and the prediction error encoding for the brain regions indicated on the top. Dots represents individual participants. Shaded areas indicate 95% confidence intervals. Dotted regression lines indicates non-significant results, while solid lines significant ones.

The above results unravel both, a neural signature of how the brain adapted its sensory representations to the predictability of inputs and how the brain encoded prediction errors. This allowed to test our key hypothesis, i.e. that, during learning, stronger encoding of prediction errors is associated with stronger representational shifts. We focused our analysis on those two clusters of brain regions that showed a significantly stronger representational shift for the high as compared to low regularity sequence: left temporal cortex and bilateral frontal cortices. Indeed, we found a significant positive correlation (Fig. 3D) between the magnitude of the prediction error and the representational shift in left temporal cortex across subjects (r = 0.47, p = 0.021 Bonferroni-corrected). Frontal cortices showed no significant effect (r = 0.22, p = 0.287 Bonferroni-corrected). In sum, in accordance with our central hypothesis, we found that the stronger the prediction error signal was encoded in left temporal cortex the stronger was the updating of the representational geometry in this brain region.

We next aimed to extend our analysis framework beyond the traditional view of the brain as a collection of modular regions (41, 42). Thus, we asked if prediction error signals resulted from distributed processing across a network of brain regions rather than independent processing within each of these regions. We employed Partial Information Decomposition (33) to decompose the joint mutual information of pairs of brain regions about error signals 90-120 ms post tone onset into redundant and synergistic components (43, 44) (Fig. 4A). This allowed us to investigate if networks of brain regions processed the prediction error either in a similar, but independent way (redundancy) or in a complementary, distributed fashion (synergy) (45).

**Figure 4:**
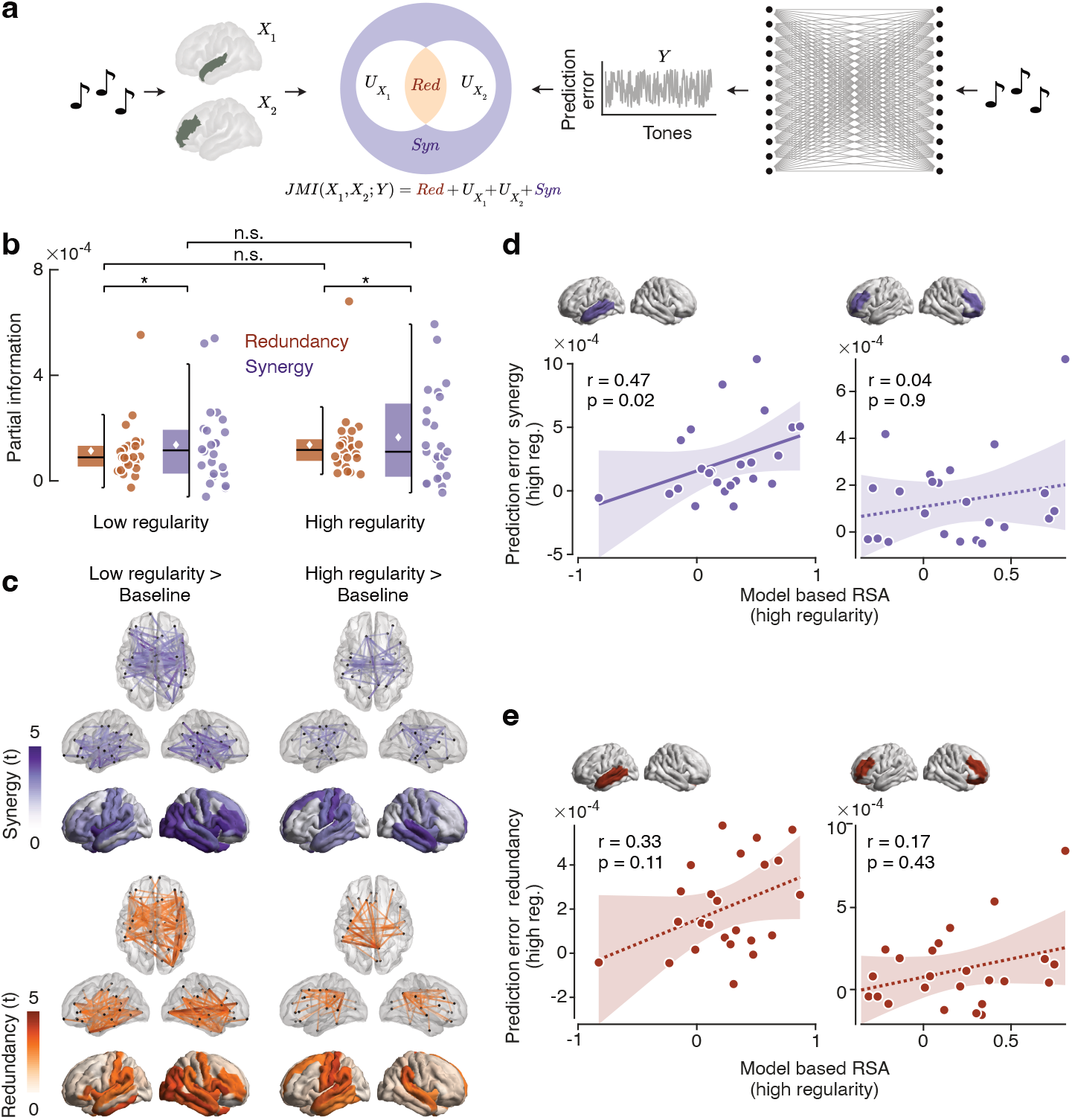
Partial Information Decomposition (PID) of the joint mutual information between pairs of brain areas encoding prediction errors. **a**, Analysis framework for computing PID from the pairwise joint mutual information (center), where the predictors (*X*_1_ and *X*_2_) are the brain signal across trials at the time window where the prediction error was mostly encoded (90-120 ms, left), and the predicted variable is the prediction error trajectory (*Y*, right). **b**, average redundancy and synergy of prediction error encoding across all pairs of brain areas. Dots represent participants and asterisks indicate statistical significance (p < 0.05). **c**, Redundancy and synergy t-statistics against the pre-tone baseline. Top views show significant cortical interactions. Bottom views show the betweenness centrality of each brain region. Transparency indicates statistical significance (p < 0.05 corrected). **d**, correlation analysis between the representational shift and betweenness centrality of the synergy component for the brain regions indicated on the top. Dots represents individual participants. Shaded areas indicate 95% confidence intervals. Dotted regression lines indicates non-significant results, while solid lines significant ones. **e**, correlation analysis between the representational shift and betweenness centrality of the redundancy component.

We found that for both, high and low regularity conditions and across all pairwise brain regions the neural interactions encoding the prediction error were substantially synergistic, and this synergy was even significantly higher than the redundancy component (low regularity: p = 0.035, d = 0.44; high regularity: p = 0.038, d = 0.43). There was no significant difference of redundancy (p = 0.549) or synergy (p = 0.502) between low and high regularity conditions. To pinpoint which cortical interactions involved redundant and synergistic encoding, we contrasted these information components between the interval with strongest prediction error encoding (90-120 ms post tone onset) and the pre-tone baseline (-50 to 0 ms) using network-based statistics (NBS) (46). For both information components and regularity conditions, we found a large-scale network of interactions, involving mostly frontal, parietal, and temporal areas (Fig. 4C, views with connections) (all p < 0.05 component-corrected). Also, the betweenness centrality of synergy and redundancy, revealed frontoparietal and temporal cortices as hubs for both information components (Fig. 4C, views with shaded regions, all p < 0.05 cluster-corrected) with no significant difference between regularity conditions (interactions: all p > 0.05 component-corrected; betweenness centrality: all p > 0.05 cluster-corrected). In sum, we found that the encoding of prediction errors involved not only redundant but also synergistic interactions across a frontoparietal and temporal network.

Do these synergistic interactions predict the representational shift during statistical learning? To address this, we repeated the same correlation analysis on the selected clusters that we performed on the ‘independent’ prediction error encoding and representational shift, but this time using the betweenness centrality of prediction error encoding synergy and redundancy (Fig. 4D-E). Indeed, we found that the centrality of synergy of the left temporal cortex significantly correlated with the representational shift across participants (Fig. 4D, r = 0.47, p = 0.022 Bonferroni-corrected), while in bilateral frontal cortices this was not the case (r = 0.03, p = 0.902 Bonferroni-corrected). For redundancy (Fig. 4E), neither left temporal cortex (r = 0.33, p = 0.113 Bonferroni-corrected) nor bilateral frontal cortices (r = 0.17, p = 0.434 Bonferroni-corrected) showed a significant correlation. In sum, we found that the temporal cortex synergistically encoded prediction errors with a large-scale network of brain regions, and that this synergistic encoding predicted the updating of sensory representations during learning.

## Discussion

Our results provide new insights on how prediction error signals shape the representational geometry of the human brain.

Our findings indicate a representational shift, where the sensory representations of contiguous and predictable tones became more similar. This effect is commonly referred to as chunking (47). Our results accord well with a large number of studies that provided indirect evidence for chunking (20, 34, 48, 49, 49–53) as well as few previous studies that directly reported neural representations to be chunked according to the predictive structure of a sensory sequence (26, 28). Our result extend these findings based on a temporally resolved RSA (29) which allowed us to track the representational dynamics throughout learning. In line with previous work (18, 26, 28), our results revealed changes of the representational geometry consistent with chunking in both sensory and high-level brain regions. This suggests distinct computational systems across the cortical hierarchy tracking sensory statistics in parallel, possibly coordinated by hippocampal activity (28, 54). Notably, the temporally resolved RSA also allowed us to track the representational dynamics on a fast timescale for each tone presentation. This revealed an early latency of the chunking effect around 120 ms post tone-onset, which temporally overlaps with the peak pitch representation. This suggests that once representational changes are established, they do not require further top-down modulation (55). It is important to note that this effect may reflect the naturalistic and continuous presentation of stimuli in our paradigm. Different outcomes may arise in segmented, trial-based settings.

Similar to the neural processing sequences shown here, also the dynamics of deep neural networks display a chunking effect during training when gradient-based methods guide them in classification tasks (56, 57). Initially random in high-dimensional space, the hidden layers’ representations become organized to distinguish class instances effectively through learning. This process mirrors the brain’s processing strategy and suggests shared computational principles between natural and artificial systems (58–60). Moreover, both systems might cluster similar sensory data to optimize energy use, adhering to environmental and computational limits, thus minimizing extraneous exploration of the sensory state space in favor of more streamlined information processing (61–64).

Our results also shed light on the neural mechanisms underlying the cortical encoding of prediction errors. We exposed an ideal observer model (30, 31, 39, 65) to the same sequences seen by human participants to extract theoretical precision-weighted prediction error trajectories. This computational approach allowed us to study prediction error encoding beyond the traditional comparison of standard and deviant stimuli in oddball paradigms (10, 11, 15, 48) in a more naturalistic sequence paradigm (66). Prediction error encoding peaked around 100 ms after each tone was delivered. As for the temporal dynamics of the representational shift, this relatively early latency in comparison to evidence reported on the mismatch negativity (10, 15) could be ascribed to the specific experimental paradigm involving a continuous stimulus presentation. Also, the latency of the prediction error encoding temporally overlapped with pitch encoding. This accords well with recent evidence for the encoding of pitch and pitch expectations at similar latencies and different cortical sites in the human auditory cortex (67).

We found that error signals were encoded in a large-scale network involving temporal and frontoparietal cortices. Thus, in line with previous evidence, error signals were encoded at both sensory and high-level processing stages (31, 68–70). Importantly, we found no difference in prediction error encoding between low and high regularity conditions, suggesting that these brain areas encode the error signal in a context-independent manner, regardless of the volatility of environmental statistics (71, 72).

The large-scale network of prediction error encoding led us to investigate how neural interactions between brain regions (73–75) contribute to error encoding. There is increasing evidence suggesting that cognitive and sensory processing in the brain is carried out in a distributed fashion rather than being localized (74, 76–78) or, in other words, that the whole is greater than the sum of its parts (44). To address this, we leveraged Partial Information Decomposition (PID) (33), which allows to decomposed the information dynamics between brain regions into synergistic and redundant interactions (45). Here, redundancy implies that two brain regions encode the prediction error in the same way, indicating a common encoding mechanism (33, 45, 79). In contrast, synergy reflects the tendency of the two brain regions to complementarily encode the error signal, indicating a distributed encoding mechanism (33, 44, 45). We found that, on average, the neural interactions encoding prediction errors were synergistic rather than redundant, suggesting a distributed rather than a shared encoding mechanism (33, 44, 45). These findings, to our knowledge, provide the first evidence for synergistic interactions underlying the encoding of prediction errors in the human brain. These results add to a growing body of evidence that challenges the traditional view of the brain as a collection of modular areas (41, 42) and suggests that predictive cortical processing is distributed rather than localized (44, 74, 76–78). Furthermore, our results suggest that the dominance of synergistic encoding in the human brain is independent of contextual regularity. This is consistent with recent evidence from the marmoset brain suggesting that predictive processing is characterized by synergistic dynamics (43).

When we inspected neural interactions both at the network level and among the cortical hubs accounting for most of the information dynamics, we found that fronto-parietal and temporal regions were strongly interrelated by both redundant and synergistic interactions. Some of the strongest synergistic interactions involved the left and right auditory cortices, indicating that the two hemispheres can integrate information in a complementary fashion. Such pairwise interactions could be mediated by higher-order interactions involving other brain regions (80, 81). Importantly, the cortical distribution of synergistic encoding strongly overlapped with the results of modular searchlight analysis. Furthermore, the same areas that showed the correlation between error computation and representational shift were also cortical hubs of the synergistic interactions broadcasting the error signal throughout the cortex. This further supports the idea that prediction error encoding results from a network computation rather than local processing (41, 82, 83).

These findings posit possible challenges for the hypothesized hierarchical nature of predictive processing in some theories of predictive coding (6, 19, 21, 70, 84). For example, in the Rao and Ballard model (6) the prediction error is computed independently at each hierarchical level. However, the large-scale synergistic component in our results suggests that each brain region encodes only partial information needed for the error computation, which is then broadcasted into the network and integrated with feedback and recurrent connections to eventually process the prediction error (43). Thus, our results support theories of predictive processing that do not necessarily require the hierarchical processing postulated by traditional predictive coding theories of perception (85).

Besides constraining the architectural aspects of predictive processing, our results provide critical evidence for the core hypothesis of the predictive coding framework, i.e. that the brain employs a generative model of the world and uses prediction errors to update model representations (4–6). To our knowledge, our results provide the first direct evidence linking these two fundamental information processing primitives, prediction error encoding and representational change. We found that the brain areas that manifested a strong representational shift, which at the same time was predicted by synergistic prediction error encoding corresponded to sensory areas such as the left auditory cortex. This may reflect a particular sensitivity of sensory areas to the modulation of their representational content (86, 87). At the same time, these areas could be an important target for top-down signals required for comparing sensory expectations and observations.

In conclusion, our findings provide evidence that largescale neural interactions engaged in predictive processing modulate the representational content of sensory areas, which may enhance the efficiency of perceptual processing in response to the statistical regularities of the environment.

## Acknowledgements

This study was supported by the European Research Council (ERC; https://erc.europa.eu/) CoG 864491 (M.S), by the German Research Foundation (DFG; https://www.dfg.de/) projects 276693517 (SFB 1233) (M.S.) and SI 1332/6-1 (SPP 2041) (M.S.), and by the German Federal Ministry of Education and Research (BMBF) to the German Center for Diabetes Research (DZD01GI0925, H.P.).

## Author contributions

AG: Conceptualization, Software, Formal analysis, Visualization, Writing – original draft, Writing – Review & Editing

JM: Methodology, Investigation, Data curation, Software, Writing – Review & Editing

HP: Methodology, Supervision, Writing – Review & Editing

MS: Conceptualization, Supervision, Resources, Project administration, Funding acquisition, Writing – original draft, Writing – Review & Editing

## Data availability statement

Raw MEG data to reproduce all the results in our study are openly available at Zenodo via the following link: http://doi.org/10.5281/zenodo.3961467.

## Materials and Methods

### Participants

Participants were 24 healthy volunteers (12 male) between 20 and 37 years old (mean age 27.54 years, SD = 9.96). All participants were right-handed and had normal hearing abilities. The experiment was realized in accordance with the Helsinki Declaration and the local ethics committee of the Medical Faculty of the University of Tübingen approved the study (No. 231/2018BO1). Data were previously published in Moser et al. (34).

### Stimuli

Stimuli consisted of 12 pure sinusoidal tones between 261.63 and 932.33 Hz (Fig. 1A). The 12 tones coincided to the musical notes C, D, E, F#, G# and A# from the 4th and 5th octave of a standard piano (261.63 Hz, 293.66 Hz, 329.63 Hz, 369.99 Hz, 415.3 Hz, 466.16 Hz, 523.25 Hz, 587.33 Hz, 659.26 Hz, 739.99 Hz, 830.61 Hz, 932.33 Hz). These tones served to create two sequences with different regularities. Both sequences consisted of 2400 tones (Fig. 1B, lasting 300 ms and presented every 333 ms) clustered into triplets (lasting 1 s) composed of three tones that never spanned more than one octave (34). In each sequence, neither the same tone, nor the same triplet of tones, could repeat consecutively twice. In the high regularity sequence (Fig. 1C), there were only four types of triplets repeating over the course of the sequence. The order of tones inside a triplet was counterbalanced across participants, yielding three different combinations of the high regularity sequence. In the low regularity sequence (Fig. 1C), each triplet changed constantly throughout the sequence, albeit maintaining the octave constrain.

### Procedure

After completing a short hearing assessment with a screening audiometer (Hortmann Neuro-Otometrie Selector 20 K) to confirm normal hearing, participants seated in a height-adjustable chair, inside a magnetically shielded room, and were told to fixate on a cross during the course of the experiment. Auditory stimulation was presented through earplugs at an intensity of 70 dB. Participants were instructed to passively listen to the sounds with no particular task to perform. The order of conditions was not counterbalanced, as the high regularity sequence always followed the low regularity sequence, after a short break, for all participants (34).

### MEG data acquisition and pre-processing

MEG data were recorded using a 275-sensor, whole-head CTF MEG system (VSM Medtech, Port Coquitlam, Canada) installed in a magnetically shielded room (Vakuumschmelze, Hanau, Germany). The sampling rate MEG signal was 585.94 Hz. We applied a fourth-order butterworth bandpass filter (0.5 to 40 Hz) and epoched the data by firstly correcting the trigger channel for a 32 ms sound output delay (34), and then selecting all the tones from 0 to 330 ms. Next, we resampled the data to 200 Hz and rejected noisy channels using a semi-automatic procedure, involving visual inspection and a cutoff threshold of root mean square (RMS) > 0.5 pT. We applied Independent Component Analysis (ICA) (88) to decompose the signal and discard eye movement, muscular and cardiac artifacts, using FastICA (89) with the number of independent components reduced to 50. The estimated independent components were visually inspected and rejected based on their topological, temporal and spectral characteristics whenever they showed an artifactual profile (90).

### Source reconstruction

After pre-processing, MEG sensors were aligned to the brain template “fsaverage” (91), from which we generated a single shell head model to compute the physical forward model (92) using FieldTrip (93). Source coordinates, head model and MEG channels were co-registered on the basis of the nasion, left and right preauricular points. We used sensor-level MEG data, aggregated from both conditions, to estimate the filter weights of a Linearly Constrained Minimum Variance (LCMV) beamformer (36), with the regularization parameter set to 5%. This spatial filtering approach reconstructs source activity with unit gain while, at the same time, maximizing the suppression of contributions from other neural sources (36). We fixed the orientation of the dipoles using singular value decomposition (SVD) to pick the direction that maximized the power (94). Then, we used the filter weights to project single-trial MEG sensor data to the source space, correcting for the sign-flip due to the SVD applied for selecting the optimal orientation that maximizes output power. The source space was finally parcelled into 72 different brain areas using the Desikan-Killany parcellation scheme (35).

### Representational Similarity Analysis

We firstly analyzed source-reconstructed MEG using a Representational Similarity Analysis (RSA) approach. We split the trials (each tone) into 5 non-overlapping blocks over the course of each sequence and computed the representational dissimilarity matrices (RDMs) at each block separately (Fig. 2A). We used the cross-validated Mahalanobis distance (cvMD) (37, 38) as a dissimilarity metric with a 10-fold cross validation scheme, due to its renowned statistical properties especially suited for RSA with neuroimaging data (37, 95). We applied the Ledoit-Wolf method (96) to compute the asymptotically optimal shrinkage parameter to regularize the covariance matrix from the training set. We ordered the RDMs entries in both sequences to have the diagonal representing the distance between tones belonging to each triplet in the high regularity sequence (Fig. 2A).The rows of the RDMs corresponded to the first, second and third tone in each triplet, ordered from the first to the fourth triplet (e.g., the fourth row was the first tone of the second triplet, which could have been either tone *D*_4_, *G*#_4_ or *C*_5_). RDMs were computed either in a time-resolved fashion (97), using all brain parcels as features for each time point, or in a searchlight manner (98), using as feature each brain parcel alongside its 5 spatial nearest neighbours for a certain time window (averaging across the time dimension). Then we averaged the values on the diagonal to investigate how, through learning, the tones within a triplet distanced between each other, as well as off the diagonal to study the same effect between tones that were not belonging to the same triplet. To summarize this pattern, we adopted a model-based RSA (99) by designing a RDM which had zeros on the on-diagonal entries and ones on the off-diagonals. We fitted this model-based RDM to each brain-derived RDM using Spearman correlation. Finally, we computed the slope of the on-diagonal, off-diagonal and model-based estimates across the blocks using the ordinary least square estimator with a counting vector increasing from 1 to 5 as regressor.

### Ideal observer model

We fitted prediction error trajectories extracted from an ideal observer model to the source-reconstructed MEG data. The employed ideal observer model can be conceived as a perceptron neural network (Fig. 3A), receiving as input one tone at a time and attempting to predict the next one. We represented the stimuli categorically as a one-hot encoded vector of the same length as the number of different tones used in the acoustic sequences *x*_*t*_ ∈ ℝ_1×*n*_, where *n* is equal to 12 and *t* indexes the tones in a sequence. Therefore, the input layer had the same dimensionality as the output layer. The trainable model parameters were encoded in a weight matrix *W*_*t*_ ∈ ℝ_*n*×*n*_, which connected the input and output layer. The weight matrix was initialized as a uniform prior over the categorical distribution of the tones, with all values having *n*_−1_ as entries. Given each tone, the model predicted the next one according to the following equation:

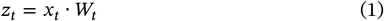

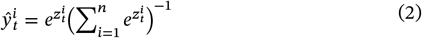

where 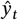 represents the prediction of the model for the next tone. We defined the loss function *ℒ* as the maximum likelihood or cross-entropy function:

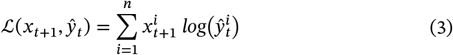

The model was trained by passing all the tones from one sequence at a time and after each observation, we computed the partial derivative of the loss function with respect to *W*_*\*t*_:

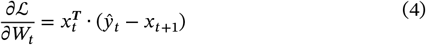

This gradient, combined to a dynamic learning rate parameter, gave rise to our measure of prediction error:

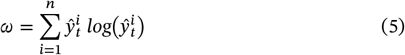

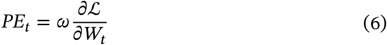

Here, the learning rate ω is not fixed as in classical reinforcement learning models (30) but depends on the uncertainty of the model since it is the Shannon entropy of the predictive distribution. This precision-weighted prediction error can account learning phenomena better than classical models with fixed learning rate (39, 100). Finally, we updated the parameters *W*_*t*_ using the gradient descent algorithm:

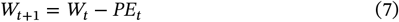

This model can also be viewed as a categorical and dynamic version of the Rescorla-Wagner model (30). We extracted the trajectory of the prediction errors, separately for each sequence, and fitted it to each parcel of the brain, for each participant. To fit the prediction error trajectories, we adopted the Gaussian Copula Mutual Information (GCMI) method (Fig. 3A), a robust multivariate statistical framework that combines the statistical theory of copulas with the analytical solution for the Shannon entropy computation of Gaussian variables (40). We first transformed each variable (brain data and prediction error trajectories) into a Gaussian variable using the inverse normal transformation. For each variable under consideration, the transformed value was obtained as the inverse standard normal cumulative distribution function (CDF) evaluated at the empirical CDF value of that variable (40). After this procedure, the mutual information is computed parametrically for Gaussian variables as following:

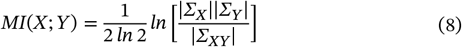

where Σ_*X*_ and Σ_*Y*_ are the covariance matrices of *X* and *Y*, respectively and Σ_*XY*_ is the covariance matrix for the joint variable (*X, Y*). In our study, we considered *Y* always as the (univariate) prediction error trajectory and *X* as the multivariate brain data. The inverse normal transformation for the brain data was applied to each feature univariately. GCMI values were computed either in a time-resolved fashion (97), using all brain parcels as variables for each time point, or in a searchlight manner (98), using as *X* each brain parcel alongside its 5 spatial nearest neighbours for a certain time window (averaging across the time dimension). Finally, we baseline-corrected these values by subtracting the GCMI values computed before the onset of the stimulus *x*_*t*_ in the time window from -50 ms to 0 ms, to avoid possible confounds given by the autocorrelation function of the two sequences.

### Partial Information Decomposition

We analyzed the MEG data using an information-theoretic approach, to investigate how brain areas interact when jointly encoding the prediction error signal. We employed Partial Information Decomposition (PID) (33) to decompose the joint mutual information (JMI), which is the information that two variables *X*_1_ and *X*_2_ give about a third target variable *Y*, in terms of different kinds of informational atoms:

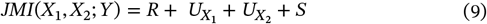

information provided by one variable but not the other (denoted as unique information, 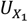 or 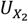), information provided by both variables separately (redundant information, *R*), or jointly by their combination (synergistic information, *S*). In this study, we considered *Y* always as the prediction error trajectory and *X*_1_ and *X*_2_ as pairs of brain parcels, for each time point in certain time window. Thus, following the intuition of the PID framework (33), we computed the redundancy measure as the minimum intersection in the information (*I*_*min*_) provided by both *X*_1_ and *X*_2_ about *Y* as follows:

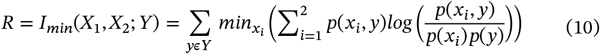

Finally, all the remaining terms can be computed using linear algebra as follows:

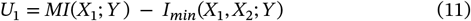

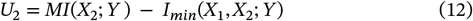

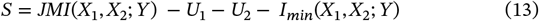

All these quantities were computed by firstly normalizing the variables using the same procedure as above for the GCMI (i.e., the inverse normal transformation). This allowed us to compute a closed form of the quantities following a parametric Gaussian model (40, 101). We computed PID components for all combinations of two brain parcels and extracted only the redundancy and synergy terms, separately for each sequence. This procedure yielded two adjacency matrices of 72 × 72 representing the pairwise neural interactions encoding the prediction error, one for the redundant and the other for the synergistic interactions. Again, we baseline-corrected these values by subtracting the redundancy and synergy computed in the baseline. Then, we averaged the matrices as a measure of global efficiency of the redundant and synergistic network, thus yielding one value per participant and sequence. Finally, we also computed the betweenness centrality measure as a measure of node importance by marginalizing the redundancy and synergy values across the interaction dimension.

### Statistical analysis

All statistical analyses were carried out at the group level (random effects) using mass univariate cluster-based paired two-tailed permutation t-tests (102) with a significance threshold α set to 0.05, 10 000 iterations, maxsum as cluster statistic and the topological neighbourhood structure defined by the proximity of the brain parcels. For the grand average redundancy and synergy, we carried out paired two-tailed t-tests, while for the statistical comparison of adjacency matrices we used network-based statistics (NBS) (46) with a significance threshold α set to 0.05, 10 000 iterations, and the size of the connected component as component statistic. For the correlation analyses, we computed the right-tail Pearson’s correlation coefficient for each selected brain cluster between the representational shift effect and the encoding prediction error effect (GCMI, redundancy and synergy centrality) across participant and correcting the multiplicity by using the Bonferroni correction. The brain clusters were selected using the first step of the cluster-based permutation test used above, i.e. by selecting the contiguous brain parcels who surpassed the alpha threshold set as above (0.05).

## References

1. Perruchet, P. & Pacton, S. Implicit learning and statistical learning: one phenomenon, two approaches. Trends in Cognitive Sciences 10, 233–238 (2006).

2. Schapiro, A. & Turk-Browne, N. Statistical Learning. In Brain Mapping (ed. Toga, A. W.) 501–506 (Academic Press, Waltham, 2015). doi:10.1016/B978-0-12-397025-1.00276-1.

3. Batterink, L. J., Paller, K. A. & Reber, P. J. Understanding the neural bases of implicit and statistical learning. Topics in cognitive science 11, 482–503 (2019).

4. Friston, K. A theory of cortical responses. Philosophical transactions of the Royal Society B: Biological sciences 360, 815–836 (2005).

5. Clark, A. Whatever next? Predictive brains, situated agents, and the future of cognitive science. Behavioral and brain sciences 36, 181–204 (2013).

6. Rao, R. P. & Ballard, D. H. Predictive coding in the visual cortex: a functional interpretation of some extra-classical receptive-field effects. Nature neuroscience 2, 79–87 (1999).

7. Friston, K., Kilner, J. & Harrison, L. A free energy principle for the brain. Journal of Physiology-Paris 100, 70–87 (2006).

8. Heilbron, M. & Chait, M. Great Expectations: Is there Evidence for Predictive Coding in Auditory Cortex? Neuroscience 389, 54–73 (2018).

9. Seriès, P. & Seitz, A. Learning what to expect (in visual perception). Frontiers in human neuroscience 7, 668 (2013).

10. Garrido, M. I., Kilner, J. M., Stephan, K. E. & Friston, K. J. The mismatch negativity: a review of underlying mechanisms. Clinical neurophysiology 120, 453–463 (2009).

11. Stefanics, G., Heinzle, J., Horváth, A. A. & Stephan, K. E. Visual mismatch and predictive coding: a computational single-trial ERP study. Journal of Neuroscience 38, 4020–4030 (2018).

12. St. John-Saaltink, E., Utzerath, C., Kok, P., Lau, H. C. & De Lange, F. P. Expectation suppression in early visual cortex depends on task set. PLoS One 10, e0131172 (2015).

13. Baldeweg, T. Repetition effects to sounds: evidence for predictive coding in the auditory system. Trends in cognitive sciences (2006).

14. Squires, N. K., Squires, K. C. & Hillyard, S. A. Two varieties of long-latency positive waves evoked by unpredictable auditory stimuli in man. Electroencephalography and Clinical Neurophysiology 38, 387–401 (1975).

15. Näätänen, R., Gaillard, A. W. & Mäntysalo, S. Early selective-attention effect on evoked potential reinterpreted. Acta psychologica 42, 313–329 (1978).

16. Garrido, M. I., Kilner, J. M., Kiebel, S. J. & Friston, K. J. Evoked brain responses are generated by feedback loops. Proceedings of the National Academy of Sciences 104, 20961–20966 (2007).

17. Winkler, I. Interpreting the mismatch negativity. Journal of Psychophysiology 21, 147–163 (2007).

18. Dürschmid, S. et al. Hierarchy of prediction errors for auditory events in human temporal and frontal cortex. Proceedings of the National Academy of Sciences 113, 6755–6760 (2016).

19. Wacongne, C. et al. Evidence for a hierarchy of predictions and prediction errors in human cortex. Proceedings of the National Academy of Sciences 108, 20754–20759 (2011).

20. Uhrig, L., Dehaene, S. & Jarraya, B. A Hierarchy of Responses to Auditory Regularities in the Macaque Brain. J. Neurosci. 34, 1127–1132 (2014).

21. Chao, Z. C., Takaura, K., Wang, L., Fujii, N. & Dehaene, S. Large-Scale Cortical Networks for Hierarchical Prediction and Prediction Error in the Primate Brain. Neuron 100, 1252-1266.e3 (2018).

22. Jiang, Y. et al. Constructing the hierarchy of predictive auditory sequences in the marmoset brain. eLife 11, e74653 (2022).

23. Schoups, A., Vogels, R., Qian, N. & Orban, G. Practising orientation identification improves orientation coding in V1 neurons. Nature 412, 549–553 (2001).

24. Hua, T. et al. Perceptual Learning Improves Contrast Sensitivity of V1 Neurons in Cats. Current Biology 20, 887–894 (2010).

25. Shibata, K., Watanabe, T., Sasaki, Y. & Kawato, M. Perceptual Learning Incepted by Decoded fMRI Neurofeedback Without Stimulus Presentation. Science 334, 1413–1415 (2011).

26. Schapiro, A. C., Rogers, T. T., Cordova, N. I., Turk-Browne, N. B. & Botvinick, M. M. Neural representations of events arise from temporal community structure. Nat Neurosci 16, 486–492 (2013).

27. Schapiro, A. C., Kustner, L. V. & Turk-Browne, N. B. Shaping of object representations in the human medial temporal lobe based on temporal regularities. Curr Biol 22, 1622–1627 (2012).

28. Henin, S. et al. Learning hierarchical sequence representations across human cortex and hippocampus. Sci Adv 7, eabc4530 (2021).

29. Kriegeskorte, N., Mur, M. & Bandettini, P. A. Representational similarity analysis-connecting the branches of systems neuroscience. Frontiers in systems neuroscience 4 (2008).

30. Rescorla, R. A. & Wagner, A. R. A theory of Pavlovian conditioning: Variations in the effectiveness of reinforcement and nonreinforcement. Current research and theory 64–99 (1972).

31. Barascud, N., Pearce, M. T., Griffiths, T. D., Friston, K. J. & Chait, M. Brain responses in humans reveal ideal observer-like sensitivity to complex acoustic patterns. Proceedings of the National Academy of Sciences 113, E616–E625 (2016).

32. Fritsche, M., Spaak, E. & De Lange, F. P. A Bayesian and efficient observer model explains concurrent attractive and repulsive history biases in visual perception. Elife 9, e55389 (2020).

33. Williams, P. L. & Beer, R. D. Nonnegative decomposition of multivariate information. arXiv preprint arXiv:1004.2515 (2010).

34. Moser, J. et al. Dynamics of nonlinguistic statistical learning: From neural entrainment to the emergence of explicit knowledge. NeuroImage 240, 118378 (2021).

35. Desikan, R. S. et al. An automated labeling system for subdividing the human cerebral cortex on MRI scans into gyral based regions of interest. Neuroimage 31, 968–980 (2006).

36. Van Veen, B. D., Van Drongelen, W., Yuchtman, M. & Suzuki, A. Localization of brain electrical activity via linearly constrained minimum variance spatial filtering. IEEE Transactions on biomedical engineering 44, 867–880 (1997).

37. Walther, A. et al. Reliability of dissimilarity measures for multi-voxel pattern analysis. Neuroimage 137, 188–200 (2016).

38. Nili, H. et al. A toolbox for representational similarity analysis. PLoS computational biology 10, e1003553 (2014).

39. Mathys, C. D. et al. Uncertainty in perception and the Hierarchical Gaussian Filter. Frontiers in human neuroscience 8, 825 (2014).

40. Ince, R. A. et al. A statistical framework for neuroimaging data analysis based on mutual information estimated via a gaussian copula. Human brain mapping 38, 1541–1573 (2017).

41. Thiebaut de Schotten, M. & Forkel, S. J. The emergent properties of the connected brain. Science 378, 505–510 (2022).

42. Urai, A. E., Doiron, B., Leifer, A. M. & Churchland, A. K. Large-scale neural recordings call for new insights to link brain and behavior. Nature neuroscience 25, 11–19 (2022).

43. Gelens, F. et al. Distributed representations of prediction error signals across the cortical hierarchy are synergistic. bioRxiv 2023–01 (2023).

44. Luppi, A. I. et al. A synergistic core for human brain evolution and cognition. Nature Neuroscience 25, 771–782 (2022).

45. Luppi, A. I., Rosas, F. E., Mediano, P. A. M., Menon, D. K. & Stamatakis, E. A. Information decomposition and the informational architecture of the brain. Trends in Cognitive Sciences 0, (2024).

46. Zalesky, A., Fornito, A. & Bullmore, E. T. Network-based statistic: identifying differences in brain networks. Neuroimage 53, 1197–1207 (2010).

47. Dehaene, S., Meyniel, F., Wacongne, C., Wang, L. & Pallier, C. The neural representation of sequences: from transition probabilities to algebraic patterns and linguistic trees. Neuron 88, 2–19 (2015).

48. Bekinschtein, T. A. et al. Neural signature of the conscious processing of auditory regularities. Proceedings of the National Academy of Sciences 106, 1672–1677 (2009).

49. Batterink, L. J., Mulgrew, J. & Gibbings, A. Rhythmically Modulating Neural Entrainment during Exposure to Regularities Influences Statistical Learning. Journal of Cognitive Neuroscience 36, 107–127 (2024).

50. Minier, L., Fagot, J. & Rey, A. The Temporal Dynamics of Regularity Extraction in Non-Human Primates. Cognitive Science 40, 1019–1030 (2016).

51. Saffran, J. R., Aslin, R. N. & Newport, E. L. Statistical Learning by 8-Month-Old Infants. Science 274, 1926–1928 (1996).

52. Farthouat, J. et al. Auditory Magnetoencephalographic Frequency-Tagged Responses Mirror the Ongoing Segmentation Processes Underlying Statistical Learning. Brain Topogr 30, 220–232 (2017).

53. Jin, P., Lu, Y. & Ding, N. Low-frequency neural activity reflects rule-based chunking during speech listening. eLife 9, e55613 (2020).

54. Stachenfeld, K. L., Botvinick, M. M. & Gershman, S. J. The hippocampus as a predictive map. Nat Neurosci 20, 1643–1653 (2017).

55. Demarchi, G., Sanchez, G. & Weisz, N. Automatic and feature-specific prediction-related neural activity in the human auditory system. Nat Commun 10, 3440 (2019).

56. Bengio, Y., Courville, A. & Vincent, P. Representation learning: A review and new perspectives. IEEE transactions on pattern analysis and machine intelligence 35, 1798–1828 (2013).

57. Rumelhart, D. E., Hinton, G. E. & Williams, R. J. Learning representations by back-propagating errors. Nature 323, 533–536 (1986).

58. Richards, B. A. et al. A deep learning framework for neuroscience. Nat Neurosci 22, 1761–1770 (2019).

59. Saxe, A., Nelli, S. & Summerfield, C. If deep learning is the answer, what is the question? Nat Rev Neurosci 22, 55–67 (2021).

60. Doerig, A. et al. The neuroconnectionist research programme. Nat Rev Neurosci 24, 431–450 (2023).

61. Hénaff, O. J. et al. Primary visual cortex straightens natural video trajectories. Nat Commun 12, 5982 (2021).

62. Ali, A., Ahmad, N., de Groot, E., van Gerven, M. A. J. & Kietzmann, T. C. Predictive coding is a consequence of energy efficiency in recurrent neural networks. Patterns 3, 100639 (2022).

63. Harrington, A. et al. Exploring the perceptual straightness of adversarially robust and biologically-inspired visual representations. In NeurIPS (2022).

64. Hosseini, E. & Fedorenko, E. Large language models implicitly learn to straighten neural sentence trajectories to construct a predictive representation of natural language. Advances in Neural Information Processing Systems 36, 43918–43930 (2023).

65. Meyniel, F., Maheu, M. & Dehaene, S. Human Inferences about Sequences: A Minimal Transition Probability Model. PLOS Computational Biology 12, e1005260 (2016).

66. Maheu, M., Dehaene, S. & Meyniel, F. Brain signatures of a multiscale process of sequence learning in humans. eLife 8, e41541 (2019).

67. Sankaran, N., Leonard, M. K., Theunissen, F. & Chang, E. F. Encoding of melody in the human auditory cortex. Science Advances 10, eadk0010 (2024).

68. Meyniel, F. & Dehaene, S. Brain networks for confidence weighting and hierarchical inference during probabilistic learning. Proceedings of the National Academy of Sciences 114, E3859–E3868 (2017).

69. Roumi, F. A., Planton, S., Wang, L. & Dehaene, S. Brain-imaging evidence for compression of binary sound sequences in human memory. eLife https://elifesciences.org/articles/84376/figures (2023) xdoi:10.7554/eLife.84376.

70. Parras, G. G. et al. Neurons along the auditory pathway exhibit a hierarchical organization of prediction error. Nat Commun 8, 2148 (2017).

71. Bonna, K., Hulme, O. J., Meder, D., Duch, W. & Finc, K. Brain network reconfiguration during prediction error processing. bioRxiv 2023.07.14.549018 (2023) doi:10.1101/2023.07.14.549018.

72. Hsu, Y.-F., Xu, W., Parviainen, T. & Hämäläinen, J. A. Context-dependent minimisation of prediction errors involves temporal-frontal activation. NeuroImage 207, 116355 (2020).

73. Siegel, M., Donner, T. H. & Engel, A. K. Spectral fingerprints of large-scale neuronal interactions. Nat Rev Neurosci 13, 121–134 (2012).

74. Panzeri, S., Moroni, M., Safaai, H. & Harvey, C. D. The structures and functions of correlations in neural population codes. Nature Reviews Neuroscience 23, 551–567 (2022).

75. Vinck, M. et al. Principles of large-scale neural interactions. Neuron 111, 987–1002 (2023).

76. Voitov, I. & Mrsic-Flogel, T. D. Cortical feedback loops bind distributed representations of working memory. Nature 608, 381–389 (2022).

77. Steinmetz, N. A., Zatka-Haas, P., Carandini, M. & Harris, K. D. Distributed coding of choice, action and engagement across the mouse brain. Nature 576, 266–273 (2019).

78. Breakspear, M. Dynamic models of large-scale brain activity. Nat Neurosci 20, 340–352 (2017).

79. Timme, N. M. & Lapish, C. A tutorial for information theory in neuroscience. eneuro 5, (2018).

80. Varley, T. F., Pope, M., Faskowitz, J. & Sporns, O. Multivariate information theory uncovers synergistic subsystems of the human cerebral cortex. Commun Biol 6, 1–12 (2023).

81. Varley, T. F., Pope, M., Maria Grazia Joshua & Sporns, O. Partial entropy decomposition reveals higher-order information structures in human brain activity. Proceedings of the National Academy of Sciences 120, e2300888120 (2023).

82. Siegel, M., Körding, K. P. & König, P. Integrating Top-Down and Bottom-Up Sensory Processing by Somato-Dendritic Interactions. J Comput Neurosci 8, 161–173 (2000).

83. Mesulam, M. M. From sensation to cognition. Brain 121, 1013–1052 (1998).

84. Friston, K. The free-energy principle: a unified brain theory? Nature Reviews Neuroscience 11, 127–138 (2010).

85. Hawkins, J., Ahmad, S. & Cui, Y. A Theory of How Columns in the Neocortex Enable Learning the Structure of the World. Frontiers in Neural Circuits 11, (2017).

86. Kok, P., Jehee, J. F. & De Lange, F. P. Less is more: expectation sharpens representations in the primary visual cortex. Neuron 75, 265–270 (2012).

87. Den Ouden, H. E., Kok, P. & De Lange, F. P. How prediction errors shape perception, attention, and motivation. Frontiers in psychology 3, 548 (2012).

88. Ikeda, S. & Toyama, K. Independent component analysis for noisy data—MEG data analysis. Neural Networks 13, 1063–1074 (2000).

89. Hyvarinen, A. Fast and robust fixed-point algorithms for independent component analysis. IEEE transactions on Neural Networks 10, 626– 634 (1999).

90. Hipp, J. F. & Siegel, M. Dissociating neuronal gamma-band activity from cranial and ocular muscle activity in EEG. Frontiers in human neuroscience 338 (2013).

91. Dale, R., Duran, N. D. & Morehead, J. R. Prediction during statistical learning, and implications for the implicit/explicit divide. Advances in Cognitive Psychology 8, 196 (2012).

92. Nolte, G. The magnetic lead field theorem in the quasi-static approximation and its use for magnetoencephalography forward calculation in realistic volume conductors. Physics in Medicine & Biology 48, 3637 (2003).

93. Oostenveld, R., Fries, P., Maris, E. & Schoffelen, J.-M. FieldTrip: open source software for advanced analysis of MEG, EEG, and invasive electrophysiological data. Computational intelligence and neuroscience 2011, (2011).

94. Westner, B. U. et al. A unified view on beamformers for M/EEG source reconstruction. Neuroimage 246, 118789 (2022).

95. Guggenmos, M., Sterzer, P. & Cichy, R. M. Multivariate pattern analysis for MEG: A comparison of dissimilarity measures. Neuroimage 173, 434–447 (2018).

96. Ledoit, O. & Wolf, M. A well-conditioned estimator for large-dimensional covariance matrices. Journal of multivariate analysis 88, 365– 411 (2004).

97. Cichy, R. M. & Pantazis, D. Multivariate pattern analysis of MEG and EEG: A comparison of representational structure in time and space. NeuroImage 158, 441–454 (2017).

98. Kriegeskorte, N., Goebel, R. & Bandettini, P. Information-based functional brain mapping. Proceedings of the National Academy of Sciences 103, 3863–3868 (2006).

99. Proklova, D., Kaiser, D. & Peelen, M. V. MEG sensor patterns reflect perceptual but not categorical similarity of animate and inanimate objects. NeuroImage 193, 167–177 (2019).

100. Gershman, S. J. A unifying probabilistic view of associative learning. PLoS computational biology 11, e1004567 (2015).

101. Kay, J. W. & Ince, R. A. Exact partial information decompositions for Gaussian systems based on dependency constraints. Entropy 20, 240 (2018).

102. Maris, E. & Oostenveld, R. Nonparametric statistical testing of EEG- and MEG-data. Journal of Neuroscience Methods 164, 177–190 (2007).

